# Serial quantification of brain oxygenation in acute stroke using streamlined-qBOLD

**DOI:** 10.1101/140897

**Authors:** Alan J Stone, George WJ Harston, Davide Carone, Thomas W Okell, James Kennedy, Nicholas P Blockley

## Abstract

It has been proposed that metabolic markers of baseline brain oxygenation have a role to play in the early identification of the ischemic penumbra. Streamlined-qBOLD is a magnetic resonance imaging technique that does not require exogenous contrast. It is a refinement of the quantitative BOLD methodology that provides a simplified approach to mapping and quantifying baseline brain oxygenation related parameters (reversible transverse relaxation rate (R_2_′), deoxygenated blood volume (DBV) and deoxyhaemoglobin concentration ([dHb])) in a clinically relevant manner. Streamlined-qBOLD was applied to an exploratory cohort of acute stroke patients in a serial imaging study. Detailed voxel-level analysis was used to quantify the metabolic profile of ischaemic tissue on presentation and investigate these metrics in relation to tissue outcome. Individual patient examples illustrate the appropriate interpretation of R_2_′, DBV and [dHb] in acute stroke and demonstrate the ability of this method to deliver regional information related to oxygen metabolism in the ischaemic tissue. Regional analysis confirms that R_2_′, DBV and [dHb] vary between regions of ischaemia with different tissue outcomes.

## Introduction

Ischaemic stroke is characterised by restricted blood supply to regions of tissue that may ultimately result in infarction. However, the brain can tolerate a limited reduction in perfusion if tissue oxygenation levels can be preserved. Therefore, techniques to map brain oxygenation in the acute phase of stroke may help to identify viable tissue that requires intervention to minimise the final infarct volume (Astrup et al., 1981). Positron Emission Tomography (PET) is the current benchmark for imaging oxygen metabolism in acute stroke (Ackerman et al., 1981; Baron, 1999) but this technique requires specialist expertise and equipment, and is not widely available in clinical settings.

Magnetic resonance imaging (MRI) has the potential to be a viable alternative to PET that is more widely available. Measurements related to oxygen metabolism are made possible by the inherent sensitivity of the transverse MR relaxation rate (R_2_*) to deoxyhaemoglobin. R_2_* (= R_2_ + R_2_′) is composed of the irreversible (R_2_) and reversible (R_2_′) transverse relaxation rates with respect to a spin echo. As changes in R_2_ (and hence R_2_*) are known to be affected by numerous factors aside from tissue oxygenation (An et al., 2014; 2012), R_2_′ is predicted to have better specificity to baseline brain oxygenation (Yablonskiy and Haacke, 1994). This sensitivity has previously been exploited to demonstrate that alterations in R_2_′ are dependent on the final outcome of ischaemic tissue (Bauer et al., 2014; Geisler et al., 2006; Seiler et al., 2012; Siemonsen et al., 2008; Zhang et al., 2011). However, it is well known that R_2_′ is dependent on both deoxyhaemoglobin concentration ([dHb]) and the deoxygenated blood volume (DBV) (Yablonskiy, 1998), resulting in ambiguity regarding the physiological origin of a measured R_2_′ alteration. Therefore, the ability to separate [dHb] from R_2_′ would provide a quantitative physiological metric directly related to tissue oxygenation. To achieve this, knowledge of the underlying DBV is required.

The quantitative-BOLD (qBOLD) method aims to separate out these physiological terms by modelling the transverse MR signal decay in the presence of a vascular network (An and Lin, 2000; He and Yablonskiy, 2007). These endogenous qBOLD methods are particularly suitable for application in acute stroke as they can be acquired non-invasively in a clinically relevant manner (An et al., 2015; Lee et al., 2003). However, in practice qBOLD techniques require several confounding effects to be considered including the influences of macroscopic field inhomogeneities (MFIs), underlying R_2_-weighting and off-resonance effects of partial volumes of cerebral spinal fluid (CSF). Streamlined-qBOLD (sqBOLD) is a recently proposed refinement of the qBOLD method targeted at minimising the effect of these confounds during data acquisition rather than by post-processing (Stone and Blockley, 2017). By removing these confounding effects from the acquired signal, baseline brain oxygenation maps can be acquired in a rapid acquisition (< 5 minutes), without the need for external contrast agents and using a simple analysis pipeline. As such, this approach represents a clinically practical solution for tracking the progression of oxidative metabolism in acute ischaemic stroke.

The aim of this study is to demonstrate the utility of sqBOLD for identifying regional changes in brain oxygenation during the acute phases of stroke. Here, sqBOLD is applied in a prospective cohort of patients with acute ischaemic stroke. Using detailed voxel-level analysis, the metabolic profile of ischaemic tissue is quantified on presentation and regional measures of sqBOLD parameters (R_2_′, DBV and [dHb]) are investigated in relation to tissue outcome.

## Material and Methods

### Patients

Patients with ischaemic stroke were recruited into a prospective observational cohort study regardless of age or stroke severity under research protocols agreed by the UK National Research Ethics Service committees (ref: 13/SC/0362). MRI was performed at presentation, 2 hours, 24 hours, 1 week and 1 month whenever possible. Nine consecutive patients were scanned on presentation. One patient was excluded from further analysis because the final lesion ROI could not be defined (no follow-up scan) and one patient was excluded from further analysis due to haemorrhagic transformation at the time of the presenting MRI leading to no relevant presenting data being acquired. As such, seven patients were included in the final analysis (Table 1).

**Table 1.** Patient characteristics.

| Patient | Stroke syndrome | Hemisphere | Sex | Age | NIHSS | Thrombolysed | Onset to Scan | Follow scan times |
| --- | --- | --- | --- | --- | --- | --- | --- | --- |
| P01* | PACS | Right | F | 76 | 25 | Y | 00:02:29 | 24hrs, 1wk, 1mth |
| P02* | TACS | Right | F | 90 | 19 | Y | 00:08:00 | - |
| P03 | PACS | Right | F | 69 | 6 | Y | 00:10:52 | 1wk, 1mth |
| P04 | LACS | Right | M | 76 | 10 | Y | 00:13:44 | 1wk, 1mth |
| P05 | POCS | Left | M | 68 | 3 | Y | 00:04:11 | 2hrs, 1wk, 1mth |
| P06 | TACS | Left | M | 79 | 14 | Y | 00:02:20 | 24hrs |
| P07 | POCS | Left | M | 68 | 12 | Y | 00:19:33 | 24hrs |
| P08 | PACS | Right | M | 78 | 4 | N | 01:04:19 | 24hrs (38hrs), 1mth |
| P09 | PACS | Right | M | 87 | 19 | Y | 00:13:49 | 24hrs, 1wk |
| NIHSS = National Institute of Health Stroke Scale; LACS = lacunar stroke; TACS = total anterior circulation stroke; PACS = partial anterior circulation stroke; POCS = posterior circulation stroke; NA = not available.<br>* Patient excluded from voxel-level and patient-level analysis. |  |  |  |  |  |  |  |  |

### Image acquisition

Scanning was performed on a Siemens 3T Verio scanner for all time points. Scanning protocols included diffusion weighted imaging (DWI) (three directions, 1.8 x 1.8 x 2.0 mm, field of view = 240 mm2, four averages, b = 0 and 1000 s/mm2, TR / TE = 9000 / 98 ms, 50 slices, 2 min 53 s) with apparent diffusion coefficient (ADC) calculation; T_1_-weighted MP-RAGE for structural imaging (1.8 x 1.8 x 1.0 mm, field of view = 228 mm, TR / TE = 2040 / 4.55 ms, TI = 900 ms, 192 slices, scan duration 3 min 58 s); and T_2_-weighted FLAIR turbo spin echo (1.9 x 1.9 x 2.0 mm, field of view = 240 x 217.5 mm2, TR / TE = 9000 / 96 ms, TI = 2500 ms, 58 slices, scan duration 2 min 8 s). A FLAIR-GASE acquisition was used to measure baseline brain oxygenation using the streamlined-qBOLD approach (Stone and Blockley, 2017) (96 x 96 matrix, field of view = 220 mm2, nine 5 mm slabs consisting of four 1.25 mm sub-slices, 100% partition oversampling, 1 mm slice gap, TR / TE = 3000 / 82 ms, TI_FLAIR_ = 1210 ms, ASE-sampling scheme Tstart / Tfinish / ΔT = −16 / 64 / 8 ms, scan duration 4 min 30 s). FLAIR-GASE consists of three separate components, nulling of CSF partial volumes using FLuid Attenuated Inversion Recovery (FLAIR) (Hajnal et al., 1992), minimisation of MFI using Gradient Echo Slice Excitation Profile Imaging (GESEPI) (Yang et al., 1998) and direct measurement of R_2_′ using an Asymmetric Spin Echo (ASE) (Wismer et al., 1988). This FLAIR-GESEPI-ASE (FLAIR-GASE) (Blockley and Stone, 2016) acquisition reduces confounding effects and when combined with quantitative modelling offers a streamlined qBOLD approach. For the patient presented in Figure 3, perfusion information was acquired on presentation using vessel encoded pseudo-continuous arterial spin labelling (VEPCASL) (EPI readout, 3.4 x 3.4 x 4.5 mm, field of view = 220 x 220 mm, 24 slices, TR / TE = 4080 14 ms, Labelling Duration = 1.4 s, Post-labelling Delays = 0.25, 0.5, 0.75, 1, 1.25 and 1.5 s, scan duration 5 min 55 s). Post-processing details of VEPCASL data to produce cerebral blood flow (CBF) maps have previously been described (Harston et al., 2017b; Okell et al., 2013).

### Post-processing

All image analysis was performed using the Oxford Centre for Functional MRI of the Brain (FMRIB) Software Library (FSL) (Jenkinson et al., 2012) and MATLAB (Mathworks, Natick, MA). Full details of the calculation of R_2_′, DBV and [dHb] parameter maps from the FLAIR-GASE data have previously been described (Stone and Blockley, 2017). In brief, the τ-series were motion corrected using the FSL linear motion correction tool (MCFLIRT) (Jenkinson et al., 2002) to the spin-echo image. The spin-echo image was brain extracted using the FSL brain extraction tool (BET) (Smith, 2002) to create a binary mask of brain tissue and all remaining τ-weighted volumes were brain extracted using this mask. This data was then fit on a voxel-wise basis to obtain parameter maps of R_2_′ by using a weighted log-linear fit to the mono-exponential regime (τ ≥ 16 ms (Yablonskiy and Haacke, 1994)). The intercept of this fit is in effect the log of the ASE signal at τ = 0 extrapolated from the mono-exponential regime (ln(S(τ = 0)_extrap_)). By subtracting the log of the measured spin-echo signal (ln(S(τ = 0 ms)) from this value, parameter maps of DBV can be produced, as previously described (Yablonskiy, 1998) (Equation 1).

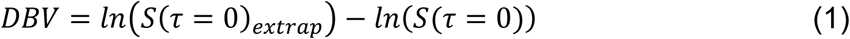

Parameter maps of [dHb] were calculated using Equation 2, where DBV and R_2_′ were measured as above and other parameters are known or assumed constants (Δχ0 = 0.264 x 10^-6^, κ = 0.03 (He and Yablonskiy, 2007)).

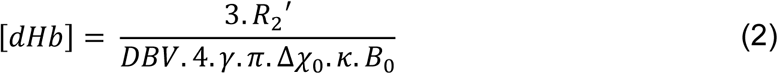

### Regions of interest

The presenting infarct region of interest (ROI) was defined using the presenting apparent diffusion coefficient (ADC) parameter map and a previously described clustering method (Harston et al., 2015), as follows. Binary masks of presenting ADC lesions were automatically generated using a threshold-defined (620 x 10-6 mm2/s) (Purushotham et al., 2015) cluster-based analysis of the ADC data. The ROI cluster was identified and smoothed (Gaussian kernel of standard deviation 1 mm) and followed by repeat cluster analysis using the FSL Cluster tool (http://fsl.fmrib.ox.ac.uk/fsl/fslwiki/Cluster). These automated ADC masks were inspected by a clinician to ensure their accuracy and manually corrected when necessary. The final infarct ROI was manually defined by an independent observer. This was done preferentially using the 1-week T_2_-FLAIR image or, if not available, the 24 hour b = 1000 s/mm2 DWI image (Harston et al., 2017a).

The following tissue outcomes were used in the analysis and were defined from the infarct ROIs in the native space of the sqBOLD parameter maps.

- The ischaemic core is tissue common to both the presenting infarct and final infarct.
- The infarct growth is tissue present in the final infarct that is not present in the presenting infarct.
- The contralateral tissue is defined by a composite mask of the presenting and final infarct tissue mirrored to the contralateral side of the brain.

### Registration

Registration of imaging modalities within a single time point was achieved using rigid body registration (6 degrees of freedom (DOF)) (Jenkinson and Smith, 2001).

Between time point registration was performed using non-linear registration of the T_1_-weighted structural scans to limit potential error introduced by edema and atrophy (Harston et al., 2017a). To create contralateral ROIs, the infarct masks were mirrored in standard space following non-linear registration of the T_1_-weighted image to a standard atlas (MNI152)(Mazziotta et al., 2001). At each time point, the FLAIR-GASE spin-echo image (τ = 0 ms) was registered (6 DOF) to the T_1_-structural using the b = 0 s/mm2 DWI image as an intermediate registration step.

### Data extraction and analysis

Voxel values of R_2_′, DBV and [dHb] were extracted from the native space of the R_2_′, DBV and [dHb] parameter maps using the ROI definitions of ischaemic core, infarct growth and contralateral tissue.

### Statistics

For each parameter (R_2_′, DBV and [dHb]), differences between the voxel-value-distributions from the tissue outcome ROIs on presentation (infarct core, infarct growth and contralateral tissue) were tested. To test the null hypothesis of no difference between the voxel-value-distributions from the tissue outcome ROIs a Kruskal-Wallis test was used. The Kruskal-Wallis test is a non-parametric version of the classical one-way ANOVA. This non-parametric test was chosen as the distribution of [dHb] values in healthy grey matter made using this technique have been shown to be better represented by the median compared to the mean (Stone and Blockley, 2017). Although the equivalent grey matter distributions of R_2_′ and DBV are normally distributed, for consistency the same non-parametric test was applied to investigate the distributions of these parameters also.

## Results

### Group characteristics

Nine consecutive large volume stroke patients were scanned on presentation, seven of which were included in the final analysis (Table 1). One patient was excluded from the analysis because the final lesion ROI could not be defined (no follow-up scan) and one patient was excluded from further analysis due to haemorrhagic transformation at the time of the presenting MRI leading to no presenting FLAIR-GASE data being acquired. The median National Institute of Health Stroke Scale at presentation was 12 (range 3 - 25) and the median symptom onset to MRI was 10 hours 52 minutes (range 2 hours 20 minutes – 1 day 4 hours 19 minutes). Eight patients received intravenous thrombolysis.

### Voxel-level analysis

To relate the presenting sqBOLD parameter maps to tissue outcome, the extracted voxel values of R_2_′, DBV and [dHb] were pooled across all patients for each tissue outcome ROI. Figure 1a-c shows distributions of parameters in each tissue type at the presenting time point and Figure 1d-f shows median and interquartile ranges of these distributions via box and whisker plots (whisker length 1.5 x interquartile range, outliers outside the whisker length are not shown). On presentation and for each parameter (R_2_′, DBV and [dHb]), the null hypothesis was rejected suggesting that the voxel-value-distributions differed significantly between the tissue outcome ROIs (Kruskal-Wallis test, p < 0.001). Following this, post hoc multiple comparisons analysis was used to perform pairwise comparisons of the three tissue outcome ROIs and revealed statistically significant differences between the tissue outcome ROIs for each parameter (infarct core and contralateral tissue; infarct growth and contralateral tissue; infarct core and infarct growth; p < 0.01 in all cases).

**Figure 1:**
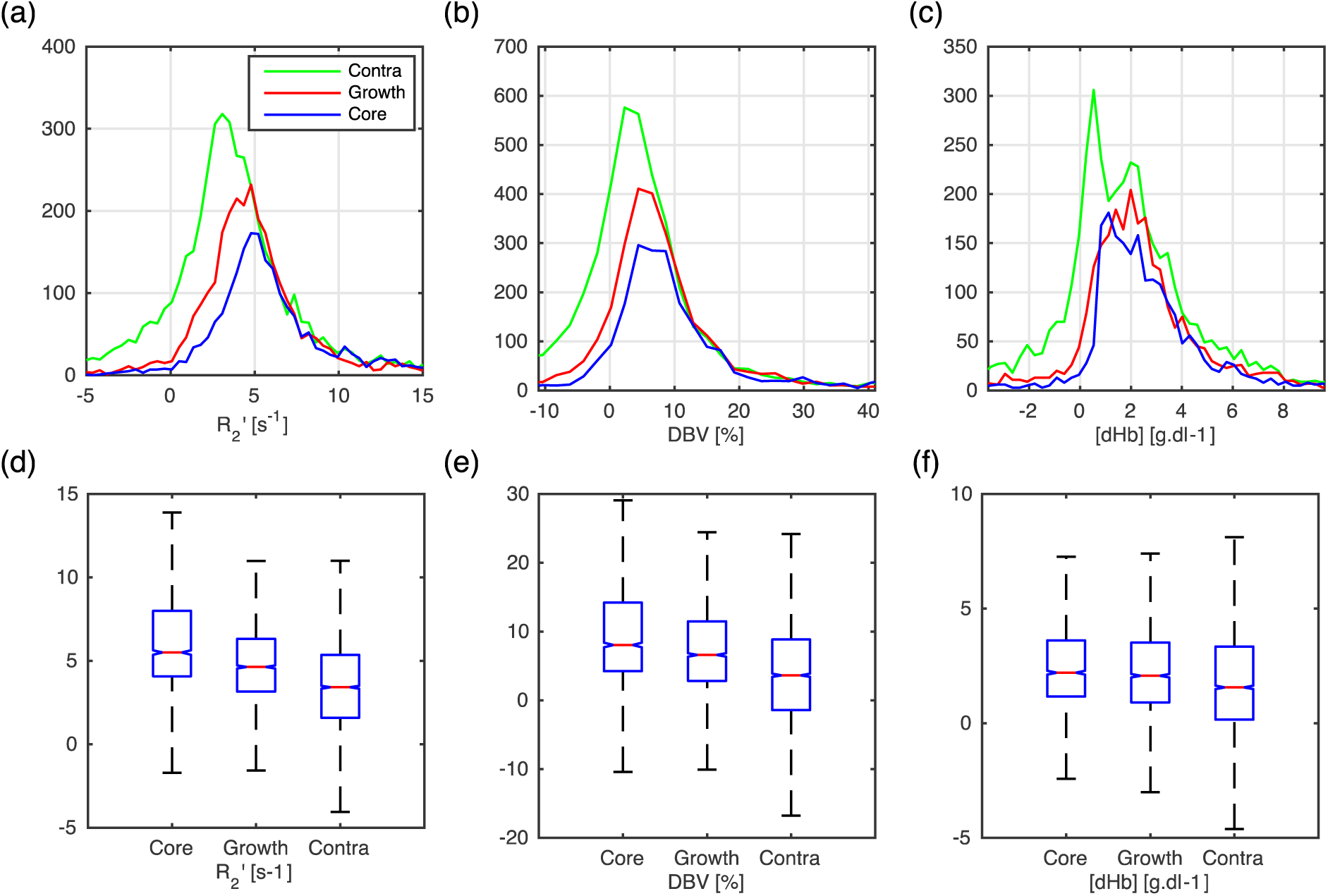
Box and Whisker plot and histograms showing voxel-level analysis of the tissue outcome from presenting time points. For each parameter (R_2_′, DBV and [dHb]) at presentation, the null hypothesis was rejected suggesting that the voxel-value-distributions differed between the tissue outcome ROIs (Kruskal-Wallis test, p < 0.001 in all cases). Post hoc multiple comparisons analysis showed statistically significant pairwise differences between the tissue outcome ROIs (infarct core and contralateral tissue; infarct growth and contralateral tissue; infarct core and infarct growth; p < 0.01 in all cases) for each parameter.

### Patient-level analysis

Using extracted voxel values, median R_2_′, DBV and [dHb] were calculated for each of the tissue outcome ROIs in each patient. Figure 2 presents group-average (± standard deviation) baseline brain oxygenation measurements for each of the tissue outcome ROIs. No significant differences were found between parameters and tissue outcome using a classical one-way ANOVA or Kruskal-Wallis test. These patient-level measures demonstrate a marked heterogeneity across the group in all parameters and this can be broadly attributed to two sources: physiological variability between patients imaged at different states of ischaemia or reperfusion and noise or error on the sqBOLD measurement. To illustrate the pertinent features of sqBOLD as applied in acute stroke, R_2_′, DBV and [dHb] parameter maps are displayed for three example patients (Figures 3 – 5).

**Figure 2:**
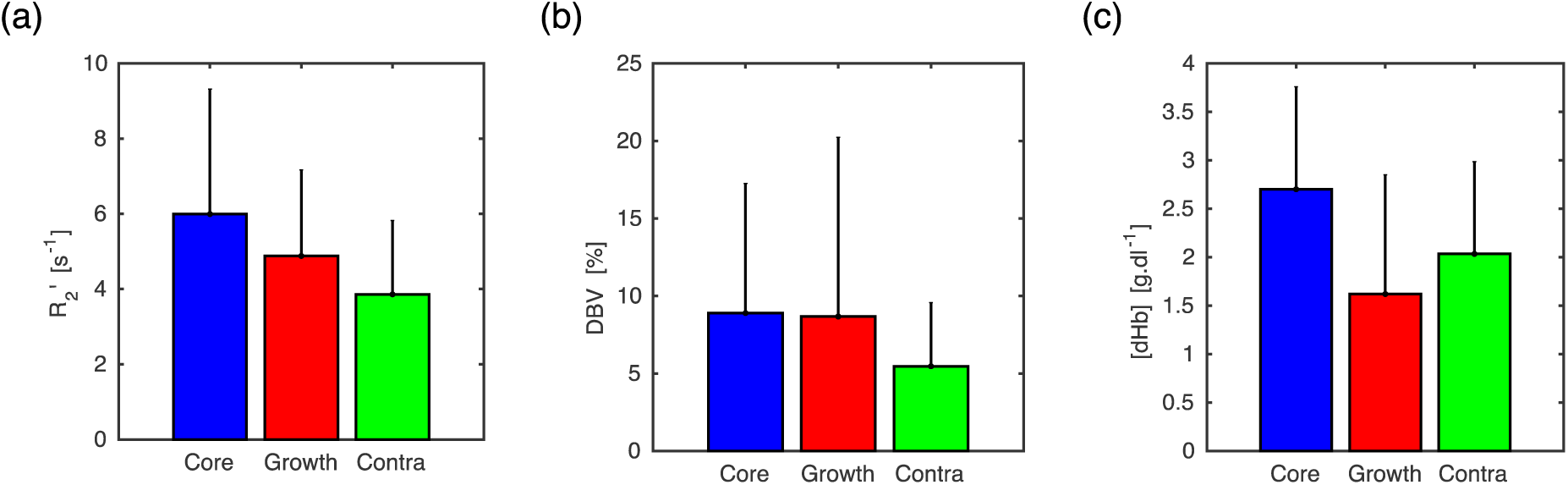
Patient-level, group average values showing tissue outcome from presenting time points. Statistical tests failed to reject the null hypothesis (p threshold = 0.05), finding no difference between the tissue outcome ROIs for any of the parameters (R_2_′, DBV and [dHb]).

**Figure 3:**
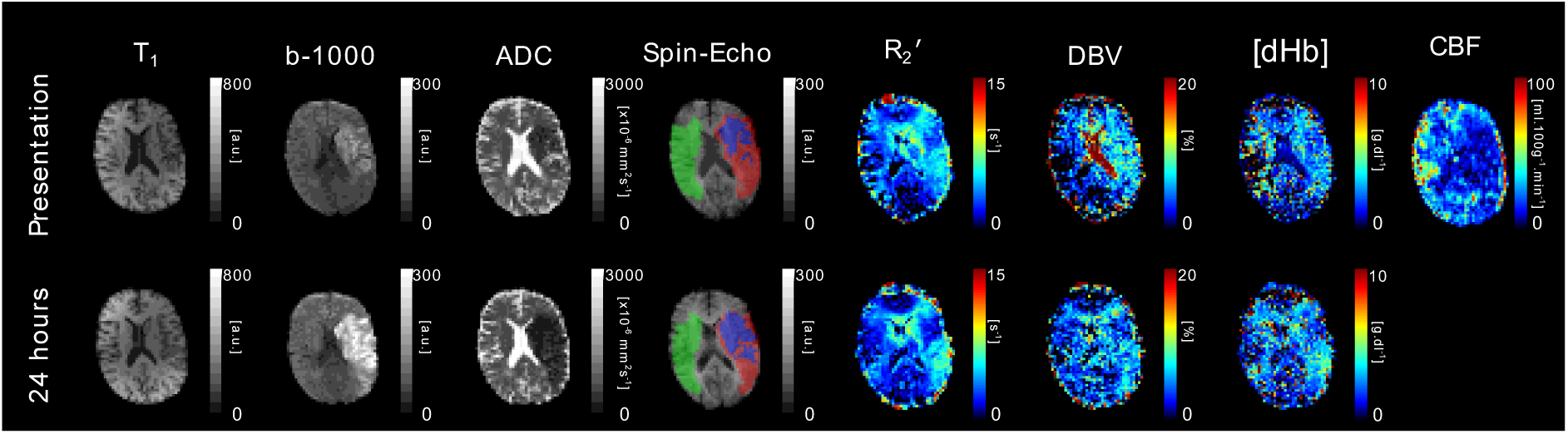
T_1_, DWI (b = 1000 s/mm^2^ and ADC) and streamlined-qBOLD parameter maps (R_2_′, DBV and [dHb]) are presented for a single axial slice in an example patient. Presenting blood flow information is displayed in the form of a CBF map. Core (blue), growth (red) and contralateral (green) tissue outcome ROIs are displayed on the spin-echo image of the streamlined-qBOLD acquisition. Patient P06 (Male, 79 years old, NIHSS = 14, thrombolysis at 1 hour 24 minutes post onset) was scanned on presentation (2 hours 20 minutes post onset) and again at 24 hours. On presentation, the DWI lesion is surrounded by areas of elevated R_2_′, DBV and [dHb]. These regions are later recruited to the 24 hour DWI lesion.

### sqBOLD parameter maps: Individual examples

For each example patient, R_2_′, DBV and [dHb] parameter maps are presented alongside T_1_-structural and DWI (b = 1000 s/mm^2^ and ADC maps) images. Ischaemic core, infarct growth and contralateral tissue ROIs are displayed on the spin-echo image of the sqBOLD acquisition.

The images in Figure 3 were acquired on presentation (2 hours 20 minutes post-onset) and 24 hours after presentation. On presentation, an apparent lesion is visible on the b = 1000 s/mm^2^ and ADC maps. The presenting R_2_′ parameter map shows a large region of elevated R_2_′ in the area surrounding the presenting DWI lesion, which goes on to infarct. At the follow up imaging time point (24 hours), the DWI lesion has grown to include the region that was elevated on the presenting R_2_′ parameter maps. The elevation in R_2_′ indicates an increase in the presence of deoxyhaemoglobin in this region which is driven by increases in DBV and/or [dHb], as seen in the accompanying parameter maps. In contrast to the large elevated region of R_2_′ there is a region of reduced R_2_′ that coincides with the presenting DWI lesion. The measure of R_2_′ in this region appears to decrease between the presenting and follow-up imaging time points.

Figure 4 shows images acquired from a second sample patient scanned on presentation (28 hours 20 minutes post onset), with follow up scanning performed at 38 hours and 1 month after presentation. A heterogeneous pattern of elevated signal can be seen on the sqBOLD parameter maps that coincide with the DWI lesion suggesting the presence of deoxyhaemoglobin in at least some regions of the diffusion lesion. In the presenting time point, apparent elevations in R_2_′ are visible in regions containing CSF. This can be ascribed to unsuccessful CSF suppression due to significant head motion.

**Figure 4:**
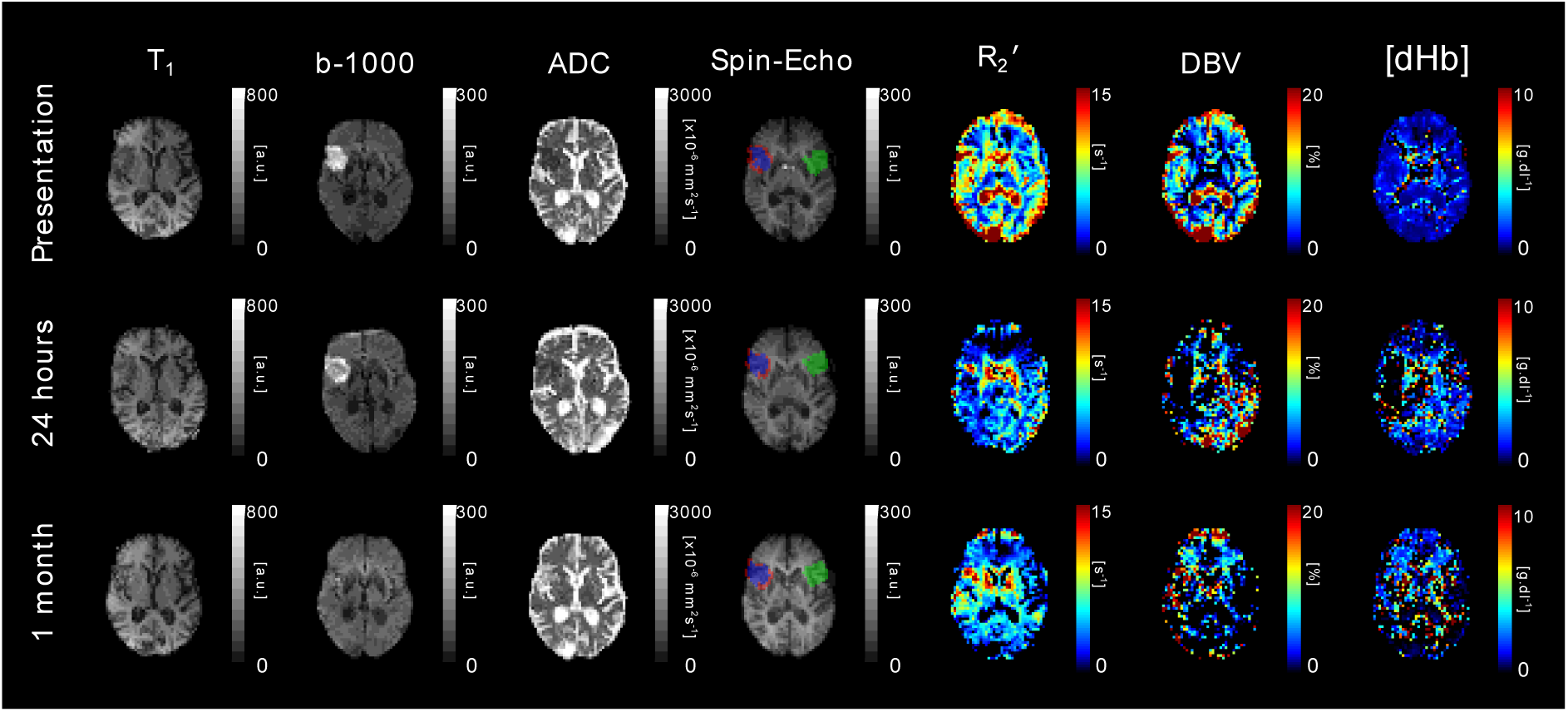
T_1_, DWI (b = 1000 s/mm^2^ and ADC) and streamlined-qBOLD parameter maps (R_2_′, DBV and [dHb]) are presented for a single axial slice in an example patient. Core (blue), growth (red) and contralateral (green) tissue outcome ROIs are displayed on the spin-echo image of the streamlined-qBOLD acquisition. Patient P08 (Male, 78 years old, NIHSS = 4, no IV thrombolysis) was scanned on presentation (28 hours 20 minutes post onset) and again at 38 hours and 1 month post initial scan. The R_2_′ maps both show a heterogeneous pattern within the DWI lesion.

Figure 5 shows images acquired from a sample patient scanned on presentation (13 hours 49 minutes post onset), with follow up scanning performed at 24 hours and 1 week after presentation. On presentation, there is an obvious deep grey matter lesion on the affected side of the brain that is clearly visible on the b = 1000 s/mm2 image. However, the presenting R_2_′ parameter maps show bilateral elevations in R_2_′ on both the affected and unaffected sides. These deep grey matter structures are known to have high iron content and the presence of this iron causes an elevation in R_2_′ that is unrelated to oxygenation and confounds the oxygenation measurement that is made within this region. This highlights the importance of considering sources of susceptibility other than deoxyhaemoglobin in the locality of the region of interest.

**Figure 5:**
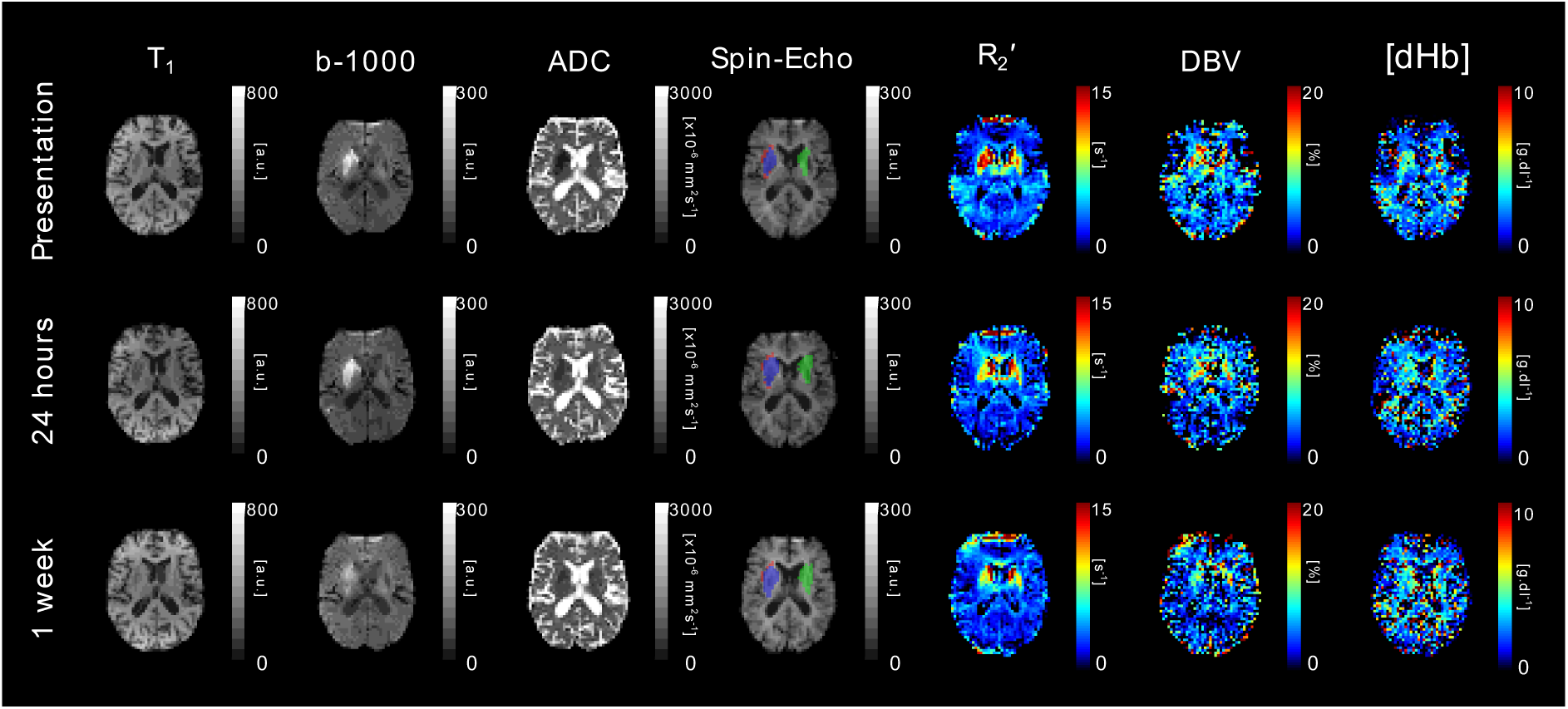
T_1_, DWI (b = 1000 s/mm^2^ and ADC) and streamlined-qBOLD parameter maps (R_2_′, DBV and [dHb]) are presented for a single axial slice in an example patient. Core (blue), growth (red) and contralateral (green) tissue outcome ROIs are displayed on the spin-echo image of the streamlined-qBOLD acquisition. Patient P09 (Male, 87 years old, NIHSS = 19, IV thrombolysis 3 hours 28 minutes post onset) was scanned on presentation (13 hours 49 minutes post onset) and again at 24 hours and 1 week post initial scan. Elevated R_2_′ matches the DWI lesion in the basal ganglia, though partly obscured by the bilateral elevation of R_2_′. [dHb] maps delineate the DWI lesion more clearly from the normal contralateral hemisphere.

## Discussion

In the acute phases of stroke, sqBOLD is shown to provide metabolic information that is indicative of the viability of ischaemic tissue. Serial-imaging from example patient cases and detailed regional analysis demonstrate that R_2_′, DBV and [dHb] are sensitive to oxygenation related changes in ischaemic tissues with varying outcomes. Significant pairwise differences in voxel distributions were observed between the regional tissue ROIs using multiple comparisons analysis for R_2_′, DBV and [dHb] (infarct core and contralateral tissue; infarct growth and contralateral tissue; infarct core and infarct growth; p < 0.01 in all cases). Median distribution values for all parameters increased in the ischaemic regions (ischaemic core and infarct growth) when compared to healthy tissue on the contralateral side (Figure 1). The most obvious increase is seen in R_2_′ and this is driven by a statistically significant increase in both deoxyhaemoglobin volume fraction (DBV) and concentration ([dHb]).

### Ischaemic penumbra

The definition of the infarct growth region used in this study is expected to be spatially and metabolically consistent with the ischaemic penumbra. In this region, an increase in [dHb] is anticipated in order to maintain the rate of oxygen metabolism in tissue that is experiencing restricted blood flow. The potential of sqBOLD to detect these changes was demonstrated by the statistically significant increase in [dHb] measured in the infarct growth ROIs across the group (Figure 1) (multiple comparisons analysis, p < 0.01). The spatial correspondence of the sqBOLD parameter maps with the infarct growth ROI can also be observed at the individual patient level **(Figure 3)**. At presentation, R_2_′, DBV and [dHb] are elevated in regions that correspond to the infarct growth ROI. The CBF parameter map, acquired in this patient on presentation, demonstrates a large region of decreased CBF that coincides with the elevated regions on the sqBOLD parameter maps. A restriction in flow is expected to result in an elevated oxygen extraction fraction (OEF), which in turn causes an increase in the relative amount of deoxyhaemoglobin produced. As such, the observation of reduced CBF and elevated [dHb] in this patient is suggestive of the early identification of tissue exhibiting the physiological traits of the ischaemic penumbra. This opens up the prospect that concurrent MR based oxygenation and flow imaging can be used to identify tissue at risk of infarction (Astrup et al., 1981). In this patient, infarction occurs in this region at some point between the presenting and follow up scan times as evidenced by the infarct growth ROI (defined from the b = 1000 s/mm2 image). Therefore, early identification of penumbral tissue would provide a window of opportunity for interventions that might salvage this tissue.

### Ischaemic core

From Figure 1, larger increases in all parameters were observed in the ischaemic core compared to infarct growth on presentation. This trend appears surprising at first, particularly if the elevated signal in the core is to be associated with the presence of deoxyhaemoglobin as a by-product of ongoing metabolism. The infarct growth region is expected to contain tissue that is metabolically active on presentation but later recruited to the final infarct volume. This is in contrast to the non-viable tissue present in the ischaemic core. However, the elevated brain oxygenation signal measured in the ischaemic core can be explained by a) stationary deoxyhaemoglobin in metabolically inactive regions with no blood supply and b) ongoing metabolism in the diffusion lesion.

### a) Stationary deoxyhaemoglobin

A similar regional trend in R_2_′ can be extrapolated from a previous study (Geisler et al., 2006) which looked at comparable tissue outcome ROIs. Here it was proposed that the elevated R_2_′ in the ischaemic core may result from stationary deoxyhaemoglobin present in vessels without blood supply. In the event of a complete occlusion of flow, stationary haemoglobin beyond the blockage will become fully deoxygenated as the remaining oxygen is metabolised leading to an increase in the amount of deoxyhaemoglobin present. This is likely to be the main contributing factor to the trend seen in Figure 1, where the ischaemic core demonstrates the largest elevation in R_2_′, DBV and [dHb]. In contrast, R_2_′ and [dHb] parameter maps in Figure 3 demonstrate a decrease in the ischaemic core. This may be explained by the presence or restoration of flow to an infarcted region. Here the metabolically inactive tissue does not produce new deoxyhaemoglobin and previously produced deoxyhaemoglobin is removed by blood flow, leading to a decrease in R_2_′, DBV and [dHb].

### b) Ongoing metabolism in the diffusion lesion

Elevated [dHb] and DBV in the ischaemic core may also be explained by observations made using PET, which have shown that regions of ongoing oxygen metabolism are possible within the presenting diffusion lesion (Fiehler et al., 2002; Guadagno et al., 2004; 2006; Kidwell et al., 2000). As such, it is possible that deoxyhaemoglobin production may still be occurring in regions of decreased ADC and contribute towards elevated R_2_′ in the ischaemic core. Parameter maps shown in Figure 4 are consistent with the heterogeneous pattern of blood oxygenation within the diffusion lesion observed using PET, but cannot be confirmed due to a lack of blood flow information.

### Interpretability of [dHb] and OEF

It is evident that the relaxometry based method used in this study is sensitive to deoxyhaemoglobin regardless of the patency of the blood supply and can therefore exhibit elevated R_2_′ in the ischaemic core. As such, knowledge of the local blood supply is important to distinguish stationary deoxyhaemoglobin present in infarcted tissue from active tissue with an elevated metabolism. This motivated the calculation of [dHb] rather than OEF to avoid the false interpretation of a high R_2_′ as elevated oxygen extraction in the absence of flow information. This sensitivity to stationary deoxyhaemoglobin also reconciles the apparent differences between PET and BOLD based measurements (Geisler et al., 2006). In PET the oxygen sensitive tracer is prevented from being delivered to the ischaemic core, meaning that signal is not detected there and reduced oxygen metabolism is inferred. This is in contrast to BOLD based measurements, which don’t rely on the arrival of a tracer and hence the presence of deoxyhaemoglobin will still cause an increase in R_2_′.

### Confounds: Non-deoxyhaemoglobin related elevations in R_2_′ and patient-motion

Streamlined-qBOLD is sensitive to other sources of susceptibility in the brain not related to deoxyhaemoglobin and care must be taken when interpreting elevations in signal. Ferritin and myelin are known sources of susceptibility that can confound the accurate quantification of brain oxygenation with this method and are of particular relevance as both can vary during ageing and in different pathologies. Figure 5 shows bilateral elevations in R_2_′ on both the affected and unaffected sides of the brain due to the high iron content of the deep grey matter structures. The presenting [dHb] map appears elevated in deep grey matter on the affected side, pointing towards the importance of interpreting the R_2_′, DBV and [dHb] parameter maps in combination, as well as being aware of non-oxygen related sources of susceptibility in the locality of the region of interest.

Significant head-motion during imaging is a challenge in acute stroke patients. Despite the segmented nature of the FLAIR-GASE acquisition, image artefacts were minimal. However, large head motions did impact the accuracy of the FLAIR CSF suppression. Slice selective inversion recovery pulses were used to optimise CSF suppression for each slice. However, large head motions between this FLAIR preparation and image acquisition resulted in ineffective removal of the CSF signal. This is evident in the presentation parameter maps in Figure 4, where elevated signal can be seen within the ventricles, particularly in the R_2_′ map. As the presence of CSF signal can lead to apparent elevations in R_2_′ not related to oxygenation, it may obscure the oxygenation changes within the diffusion lesion on presentation (Dickson et al., 2009; He and Yablonskiy, 2007; Simon et al., 2016). However, it is encouraging that heterogeneous patterns of oxygenation can be seen within the diffusion lesion at the follow-up imaging time points where CSF suppression was effective.

### Group heterogeneity, flow and further work

From the patient-level analysis (Figure 2), considerable heterogeneity was apparent across the group, meaning regional trends in R_2_′, DBV and [dHb] were not significantly different. The heterogeneity in regional parameters across the group can be attributed, at least in part, to the differences in onset to scan time (Table 1) and differences in the perfusion and reperfusion status of the ischaemic tissue. As such, it is difficult to hypothesise the expected metabolic state of this tissue on presentation, but it is likely to be varied across the group, with each patient potentially undergoing a different pathway to infarction (del Zoppo et al., 2011). This is supported by the high coefficient of variation across subjects measured in the infarct growth region, particularly for DBV and [dHb] (Figure 2).

The absence of comprehensive perfusion information is a distinct limitation of this study. Non-invasive MRI measures of blood flow can be routinely made using arterial spin labelling (ASL) (Harston et al., 2017b) and Figure 3 provides a demonstration of how flow and oxygenation measurements can be used to provide a unique insight into tissue viability. Conversely, the information provided by sqBOLD should also aid the interpretation of ASL CBF measurements, since low flow does not always progress to infarction in regions experiencing benign oligaemia (Kidwell et al., 2003). Finally, in combination CBF and [dHb] measurements would allow for the calculation of the cerebral metabolic rate of oxygen consumption (CMRO_2_) (Blockley et al., 2015). This has been shown to improve tissue outcome prediction and may partly explain the variability seen in the presenting sqBOLD oxygenation measurements (An et al., 2015).

Despite the heterogeneity of the observed signal, both between and within individuals, the pooled voxel-wise analysis (Figure 1) and the individual examples (Figures 3-5) demonstrate the potential of R_2_′, DBV and [dHb] for investigating oxidative metabolism within the ischaemic penumbra. The identification of tissue outcomes based solely on measurements from this method don’t currently appear possible, as evidenced by the considerable overlap between tissue outcome distributions in Figure 1. However, Kruskal-Wallis tests and post hoc multiple comparisons analysis found significant differences between these distributions suggesting that tissue outcome is dependent on tissue oxygenation and that the parameter maps derived from sqBOLD are sensitive to identifying this information on presentation. As such, sqBOLD provides complimentary information to existing imaging modalities such as DWI and ASL and the combination of this information may allow for earlier identification of tissue under metabolic stress during the acute phases of stroke (An et al., 2015).

In addition, this study supports the further investigation of sqBOLD in a larger scale study and highlights the importance of controlling for onset to scan time and tissue perfusion status. The non-invasive, quantitative nature of this method also means it is suitable for longitudinally monitoring stroke evolution and may provide unique insight into the various pathways to infarction and recovery, as well as providing valuable biomarkers with which to assess treatment and intervention.

## Conclusion

Streamlined-qBOLD was used to acquire information about oxidative metabolism in a cohort of acute ischaemic stroke patients, which is complimentary to conventional MRI methodologies. It was found that resting brain oxygenation related parameters (R_2_′, DBV and [dHb]) vary between regions with different tissue outcomes. The appropriate implementation of R_2_′, DBV and [dHb] parameter maps has the potential to refine the identification of the ischemic penumbra.

## Acknowledgements

This work was supported by the Engineering and Physical Sciences Research Council [grant number EP/K025716/1], the National Institute of Health Research Biomedical Research Centre Oxford, the National Institute of Health Research Clinical Research Network, the Royal Academy of Engineering, the Dunhill Medical Trust [grant number: OSRP1/1006] and the Wellcome Trust Institutional Strategic Support Fund (2014-2015). We also wish to acknowledge the staff and facilities provided by the Oxford Acute Vascular Imaging Centre.

## Appendix A. Supplementary data

The parameter maps and ROIs that underpin Figures 1-5 can be accessed via the Oxford Research Archive repository, doi: http://dx.doi.org/10.5287/bodleian:VYmwzrzpd alongside scripts which can be used to reproduce the charts in Figure 1 & 2 doi: http://dx.doi.org/10.5281/zenodo.833474

